## Supplementary Video captions for "Lomasomes and other fungal plasma membrane macroinvaginations have a tubular and lamellar genesis"

**Video S1.** Time-lapse video, *S. hirsutum* hypha fragment labeled AM4-64. It is possible to observe (arrow) the transfer of the label from the PM to the tonoplast of a large vacuole touching the PM (another variant is the division of the protoplast during plasmolysis). Observation for 30 seconds, snapping frequency 1 frame per second. Scale bar 5 µm.

**Video S2.** Time-lapse video, *S. hirsutum* mycelia labeled AM4-64. Long-term observation for 20 minutes: snapping for 3 minutes with breaks of 2 minutes, snapping frequency 1 frame per second. The formation and disappearance of PM macroinvaginations can be seen. The transverse hyphae gradually become diffusely labeled. The hypha on the right (vertical) becomes motley. Scale bar 5 µm.

**Video S3.** Time-lapse video, *S. hirsutum* mycelia labeled AM4-64. The snapping time is about 1 minute; the frequency is one frame per second. Snapping starts 5 minutes after the slide preparation. Macroinvaginations are mainly represented by glomeruli, to a lesser extent by pendants and small vesicles. White arrows indicate some of the glomeruli, yellow arrows indicate pendants. Scale bar 5 µm.

**Video S4.** Z-stack 3D-reconstruction video, *S. hirsutum* hypha fragment labeled AM4-64. It has been shown that glomeruli and very small vesicles are extended across the cell along the PM. Sinuous filaments are moving glomeruli. The arrow marks the plaque, which is also elongated across the cell. Scale bar 5 µm.

**Video S5.** Time-lapse video, *O. olearius* hypha fragment labeled AM4-64. The snapping time is about 1.5 minute; the frequency is one frame per second. The arrow indicates an invagination of the thin longitudinal tube type, which is turning across the cell and is pressed against the PM. As a result, it is visualized as a glomerulus. Scale bar 5 µm.

**Video S6.** Time-lapse video, *O. olearius* mycelia labeled AM4-64. The snapping time is about 0.5 minute; the frequency is one frame per second. The white arrow marks pendants with a small radius of movement. The red arrow indicates a local signal loss in the PM – a phenomenon not rare, but of an unknown nature. Scale bar 5 µm.

**Video S7.** Time-lapse video, *S. hirsutum* hypha fragments labeled AM4-64. A large pendant with a large radius of movement is visible (yellow arrow). Scale bar 5 µm.

**Video S8.** Time-lapse video, *O. olearius* hypha fragment labeled AM4-64. The snapping time is about 1 minute; the frequency is one frame per second. Plaques are marked with blue arrows. The yellow arrow points to a thin tube, the end of which gradually turns across the focal plane, turning into a pendant. Scale bar 5 µm.

**Video S9.** Time-lapse video, *O. olearius* hypha fragment labeled AM4-64. The snapping time is about 0.5 minute; the frequency is one frame per second. Two large pendants are visible in the center of the image. At a certain point, it becomes clear that they are small vesicles in the transverse focal section (yellow arrows). Thus, these pendants are moving thin tubes that do not immediately fall into the focal plane with their ends. Scale bar 5 µm.

**Video S10.** Z-stack video, *O. olearius* hypha fragment labeled AM4-64. The successive change of Z-stack slices shows that invaginations, which on single focal planes can be mistaken for large pendants, are transverse tubes. Scale bar 5 µm.

**Video S11.** Time-lapse video, apical hypha of *Neonothopanus nambii* labeled by AM4-64. The snapping time is about 0.5 minute; the frequency is one frame per second. The yellow arrow points to the pendant (or glomerulus) that has scissored from the PM and is carried away by the flow of the cytoplasm to the basal part of the hypha. Scale bar 5 µm.

**Video S12.** Time-lapse video, *O. olearius* mycelia labeled AM4-64. The snapping time is about 0.5 minute; the frequency is one frame per second. Yellow arrows track the movement of the labeled structure within the hyphae. Presumably, this is an invagination that has scissored from the PM from the side of the apical part of the hypha. In the last seconds of the video, it can be seen that it is a vesicle or a short tube.

**Video S13.** Time-lapse video, *S. hirsutum* mycelia labeled AM4-64. The snapping time is about 3 minutes; the frequency is one frame per second. Red arrows point to small vesicles that gradually flatten into a glomerulus and then into a small plaque. Green arrows point to the sites of forming new plaques and glomeruli. Yellow arrows mark pendants. The moving yellow arrow tracks the scission and movement of the pendant. Scale bar 5 µm.

**Video S14.** Time-lapse and Z-stack 3D-reconstruction video, *O. olearius* mycelia labeled AM4-64. The snapping time for the time-lapse mode is about 1 minute; the frequency is one frame per second. The arrow points to a large vesicle that forms from the plaque. On the right is the same location 3D-reconstructed from Z-stacks. It can be seen that the vesicle is extended across the cell. Scale bar 5 µm.

**Video S15.** Time-lapse video, *S. hirsutum* mycelia labeled AM4-64. The snapping time is about 0.5 minute; the frequency is one frame per second. Arrows point to longitudinal tubes that turn on across the cell and become visible to the observer as vesicles. The left tube is thin and turns into a small vesicle, the right tube is closer to the thick tube and turns into a large vesicle. Scale bar 5 µm.

**Video S16.** Time-lapse video, *O. olearius* mycelia labeled AM4-64. The snapping time is about 1 minute; the frequency is one frame per second. The white arrow marks two small vesicles – it can be seen that they are optical sections through two thin tubes. The red arrow points to a short, thick tube that turns on to the observer and turns into a large vesicle. Scale bar 5 µm.

**Video S17.** Z-stack 3D-reconstruction video, *S. hirsutum* hypha fragment labeled AM4-64. Some small vesicles (arrows) show that they are either short transverse tubes pressed against the PM or rolls of twisted lamellae. Scale bar 5 µm.

**Video S18.** Time-lapse video, *O. olearius* mycelia labeled AM4-64. The snapping time is about 3 minutes; the frequency is one frame per second. It can be seen how the standing thin tube grows (white arrow). At the end of the snapping, it folds or bends across the cell. The red arrow points to a thin tube growing along the PM. Scale bar 5 µm.

**Video S19.** Time-lapse video, *S. hirsutum* mycelia labeled AM4-64. The snapping time is about 3 minutes; the frequency is one frame per 5 seconds. White arrows mark growing longitudinal thin tubes. The red arrow points to a thin, standing tube that folds or bents into a large glomerulus. Scale bar 5 µm.

**Video S20.** Z-stack 3D-reconstruction video, *O. olearius* hypha fragment labeled AM4-64. Various types of tubes are presented: thin and thick, longitudinal and transverse, straight and sinuous. Scale bar 5 µm.

**Video S21.** Time-lapse video, *O. olearius* hypha fragment labeled AM4-64. Tubes of different thicknesses are presented: thin up to 500 nm (white and yellow arrows), thick over 1 µm (blue arrow) and intermediate thickness of about 750 nm (green arrow). Scale bar 5 µm.

**Video S22.** Time-lapse video, *S. hirsutum* hypha fragment labeled AM4-64. The snapping time is about 0.5 minute; the frequency is one frame per second. The video shows large vesicles. They are static. The arrow points to one of the large vesicles, inside which a concentric signal is visible - this means that the structure is elongated across the cell and narrows towards its back. Scale bar 5 µm.

**Video S23.** Time-lapse video, *S. hirsutum* mycelia labeled AM4-64. The snapping time is about 1 minutes; the frequency is one frame per second. Large vesicles have been shown to be static. The arrow points to a very large vesicle, about 3.5 µm in diameter. Scale bar 5 µm.

**Video S24.** Time-lapse video, *O. olearius* mycelia labeled AM4-64. The snapping time is about 1 minute; the frequency is one frame per second. The video shows intermediate and thick tubes. The length of the tube marked with an arrow is about 15 μm. The red arrow indicates a possible vacuole-lysosome. Scale bar 5 µm.

**Video S25.** Z-stack 3D-reconstruction video with Tikhonov-Miller deconvolution algorithm, *O. olearius* hypha fragment labeled AM4-64. A thick tube extends from the clamp-connection membrane. Scale bar 5 µm.

**Video S26.** Z-stack 3D-reconstruction video, *O. olearius* hypha fragment labeled AM4-64. Volumetric image of thick tubes. Scale bar 5 µm.

**Video S27.** Time-lapse video, *S. hirsutum* hypha fragment labeled AM4-64. The snapping time is about 1 minute; the frequency is one frame per second. In the video, it can be seen the turn of two tubes across the cell – a pseudo-transformation into the large vesicles. Scale bar 5 µm.

**Video S28.** Time-lapse video, *S. hirsutum* mycelia labeled AM4-64. The snapping time is about 1 minutes; the frequency is one frame per second. Swelling of a thick standing tube into a large vesicle is shown (arrow). Scale bar 5 µm.

**Video S29.** Time-lapse video, *S. hirsutum* mycelia labeled AM4-64. The snapping time is about 1.5 minutes; the frequency is one frame per second. Part of the thin tube pressed against the PM swells into a large vesicle. Scale bar 5 µm.

**Video S30.** Time-lapse video, *S. hirsutum* hypha fragment labeled CFDA. The snapping time is about 0.5 minute; the frequency is one frame per second. The exchange of contents between large oval vacuoles through tubular vacuoles is visible. Scale bar 5 µm.

**Video S31.** Time-lapse video, *S. hirsutum* hypha fragment labeled CFDA. The snapping time is about 1.5 minute; the frequency is one frame per second. Different types of vacuoles are presented: large oval, small round and tubular. It can be seen that tubular vacuoles are formed as outgrowths of large vacuoles, as well as by the fusion of small vacuoles. Scale bar 5 µm.

**Video S32.** Time-lapse video, *S. hirsutum* hypha fragment labeled CFDA. The snapping time is about 1 minute; the frequency is one frame per second. The video shows large strongly elongated vacuoles, but thicker than tubular vacuoles. There is a constant fusion of vacuoles with each other. These vacuoles are not visible in bright field, i.e. they are different from the large central vacuoles that are usually well visible in bright field. Scale bar 5 µm.

**Video S33.** Time-lapse video, *S. hirsutum* hypha fragment labeled CFDA. The snapping time is about 0.5 minute; the frequency is one frame per second. The hyphae are filled with many small mobile rounded vacuoles. The arrow points to tubular vacuoles connecting medium-sized rounded vacuoles. Scale bar 5 µm.

**Video S34.** Video with merged AM4-64- and CFDA-labeling, *S. hirsutum* hypha fragment. The absence of colocalization of small vacuoles and glomeruli is demonstrated. Scale bar 5 µm.

**Video S35.** Video with merged AM4-64- and CFDA-labeling, *S. hirsutum* hypha fragment. The absence of colocalization of small vacuoles and small vesicles and glomeruli is demonstrated. Scale bar 5 µm.

**Video S36.** Video with merged AM4-64- and CFDA-labeling, *S. hirsutum* hypha fragment. The absence of colocalization of large vacuoles and small vesicles and glomeruli is demonstrated. Scale bar 5 µm.

**Video S37.** Video with merged AM4-64- and CFDA-labeling, *S. hirsutum* hypha fragment. The absence of colocalization of large and small vacuoles and large vesicle (blue arrow) is demonstrated. Scale bar 5 µm.

**Video S38.** Nile Red labeling of *S. hirsutum* mycelia (red channel). Numerous small lipid droplets are visible, morphologically and dynamically different from glomeruli and pendants. Scale bar 5 µm.
